# Higher-order structures of the foot-and-mouth disease virus RNA-dependent RNA polymerase required for dynamic inter-molecular interactions involved in viral genome replication

**DOI:** 10.1101/2020.12.17.423260

**Authors:** Eleni-Anna Loundras, James Streetley, Morgan R. Herod, Rebecca F. Thompson, Mark Harris, David Bhella, Nicola J. Stonehouse

## Abstract

Replication of many positive-sense RNA viruses occurs within intracellular membrane-associated compartments. These are believed to provide a favourable environment for replication to occur, concentrating essential viral structural and non-structural components, as well as protecting these components from host-cell pathogen recognition and innate immune responses. However, the details of the molecular interactions and dynamics within these structures is very limited. One of the key components of the replication machinery is the RNA-dependent RNA polymerase, RdRp. This enzyme has been shown to form higher-order fibrils *in vitro*. Here, using the RdRp from foot-and-mouth disease virus (termed 3D^pol^), we report fibril structures, solved at ~7-9 Å resolution by cryo-EM, revealing multiple conformations of a flexible assembly. Fitting high-resolution coordinates led to the definition of potential intermolecular interactions. We employed mutagenesis using a sub-genomic replicon system to probe the importance of these interactions for replication. We use these data to propose models for the role of higher order 3D^pol^ complexes as a dynamic scaffold within which RNA replication can occur.

## Introduction

Replication of genomic material is a key stage of the lifecycle of all viruses. Some of the molecular details of replication are well defined as a result of elegant in vitro analyses. It has also been demonstrated that for many viruses, replication occurs in macromolecular complexes involving both viral and cellular components (Bienz *et al.*, 1983; Schlegel *et al.*, 1996; Gosert *et al.*, 2003; Monaghan *et al.*, 2004; Novoa *et al.*, 2005; Knox *et al.*, 2005). However, the molecular details of the architecture of replication complexes remains unknown and thus we do not fully understand how proteins and viral genetic material are associated during the replication process and any details of dynamics that are involved.

The genomes of positive-sense RNA viruses act directly as a messenger RNA and hence are translated immediately after infection. For picornaviruses such as poliovirus (PV), rhinovirus and foot-and-mouth disease virus (FMDV), proteolytic processing of the viral polyprotein ultimately releases the non-structural proteins involved in genome replication. A key protein is the RNA-dependent RNA polymerase, RdRp, termed 3D^pol^. This enzyme (in complex with other viral proteins e.g. 2C, 3A and 3CD and a replication primer protein termed 3B) produces negative strand genomic intermediates, from which new positive strands are synthesised (Flint and Ryan, 1997; Gamarnik and Andino, 1998; Goodfellow *et al.*, 2003; Nayak *et al.*, 2005).

Biochemical and structural studies have yielded a very good understanding of the steps involved in each step of nucleotide incorporation (e.g. Peersen, 2017). The structure of the picornavirus 3D^pol^ resembles a cupped right hand, in common with many other DNA and RNA polymerases (Steitz, 1998; Brautigam and Steitz, 1998). There are three defined domains, termed thumb, fingers and palm. The finger domains are extended in picornaviral 3D^pol^ structures, when compared to many other polymerases, and contact the tip of the thumb domain. This contact is important for maintaining polymerase structure whilst the geometry of the fingers against the thumb form part of the NTP entry channel. The single-stranded template RNA enters 3D^pol^ through a channel at the top of the enzyme and takes a right angle turn at the palm active site when the product-template duplex is formed. This emerges from the large front opening (Ferrer-Orta *et al.*, 2004; Ferrer-Orta, Arias, Escarmís, *et al.*, 2006; Ferrer-Orta, Arias, Agudo, *et al.*, 2006; Ferrer-Orta *et al.*, 2009; Ferrer-Orta *et al.*, 2015; Herod *et al.*, 2016). There is, however, no strand separation system identified from biochemical and/or structural analysis of the polymerase alone, but such a mechanism must presumably exist to prevent formation of a stable RNA duplex.

In contrast to its catalytic role in replication, the non-catalytic roles of 3D^pol^ are less defined. It forms part of a larger precursor 3CD, and is essential for RNA replication via genomic cyclisation (Andino *et al.*, 1990; Gamarnik and Andino, 1998; Herold and Andino, 2001). Previous studies have also suggested that 3D^pol^ (or larger precursors thereof) has an essential cis-acting role in replication which is distinct from a separate trans-acting role (Spear *et al.*, 2015; Herod *et al.*, 2016). Dimers and higher-order oligomers of 3D^pol^ molecules have been reported in a number of positive-sense RNA viruses (Hansen *et al.*, 1997; Luo *et al.*, 2000; Lyle *et al.*, 2002; Hogbom *et al.*, 2009; Chinnaswamy *et al.*, 2010;). Furthermore, the 3D^pol^ enzyme of both PV and FMDV has been previously shown to form higher-order fibrillar structures (Lyle *et al.*, 2002, Spagnolo *et al.*, 2010; Bentham et al, 2012; Tellez *et al.*, 2011; Wang *et al.*, 2013). PV 3D^pol^ has also been demonstrated to form planar lattices and higher-order structures have been visualised within infected cells (Lyle *et al.*, 2002). It has therefore been postulated that the higher-order structures act as a replication scaffold, however, no RNA has been shown within the PV 3D^pol^ structures and furthermore, these were demonstrated to form in vitro in the absence of RNA (Spagnolo *et al.*, 2010; Tellez *et al.*, 2011; Wang *et al.*, 2013). In contrast, the FMDV fibrillar structures were only detected in vitro in the presence of RNA. Indeed, our previous studies showed that primer, RNA template and nucleotides were required for fibril formation, thus suggesting that they could be part of an active replication complex (Bentham et al, 2012).

Using immunofluorescent confocal and electron microscopy, FMDV has been shown to dysregulate Golgi and ER-derived membranes in a similar way to PV (Schlegel *et al.*, 1996; O’Donnell *et al.*, 2001; Monaghan *et al.*, 2004; Knox *et al.*, 2005; Midgley *et al.*, 2013), but distinct membrane-bound replication complexes comprised of viral RNA, structural and non-structural proteins, and host-cell proteins have yet to be identified for FMDV. Although both viruses are members of the Picornaviridae, the Aphthovirus FMDV has several unique features that distinguish it from other members of the family including an extended 5’ untranslated region and triplication of the primer encoding region.

Picornaviruses are responsible for a number of important human and animal diseases. For example FMDV, the causative agent of foot-and-mouth disease, is a significant economic problem, such as occurred in the UK in 2001 (where economic loses to farming and tourism were estimated at ~ £15 billion at current value) (Thompson *et al.*, 2002; Dee *et al.*, 2018). Therefore, a more complete understanding of the composition of the replication machinery is important for therapeutic development. Here, we have trapped the products of a replication assay by cross-linking and thus captured higher-order structures in a range of conformations. Using cryo-electron microscopy, we have calculated the structures of 3D^pol^ fibrils in 9 conformations at 7-9 Å resolution, with some of these occurring within the same fibril. We also probed the importance of 3D^pol^-3D^pol^ interfaces within the fibril by mutagenesis and we use these data to build a model of dynamics of 3D^pol^ interactions in replication.

## Results

### Wild-type 3D^pol^ can catalyse incorporation of UTP into nascent RNA

We first sought to demonstrate that our recombinant 3D^pol^ protein was able to bind a synthetic template RNA and was catalytically active. We produced wild-type (WT) 3D^pol^ in *E. coli* as previously described (Ferrer-Orta *et al.*, 2004; Bentham *et al.*, 2012), alongside 3D^pol^ with mutations in the active site (3D^pol^-GNN) as a control. In order to assess the affinity of RNA binding by 3D^pol^, a fluorescence polarisation anisotropy (FPA) assay was performed. Samples of 3D^pol^-WT and 3D^pol^-GNN were titrated with FITC-labelled 13mer poly-A RNA (Fig 1A), resulting in Kd values of WT and GNN of 5.83 ± 1.3 and 1.92 ± 0.67 μM, respectively (Fig 1B). These data showed that both active and inactive enzymes were able to bind RNA *in vitro*, with the mutant demonstrating significantly higher affinity (p < 0.01).

**Figure 1.**
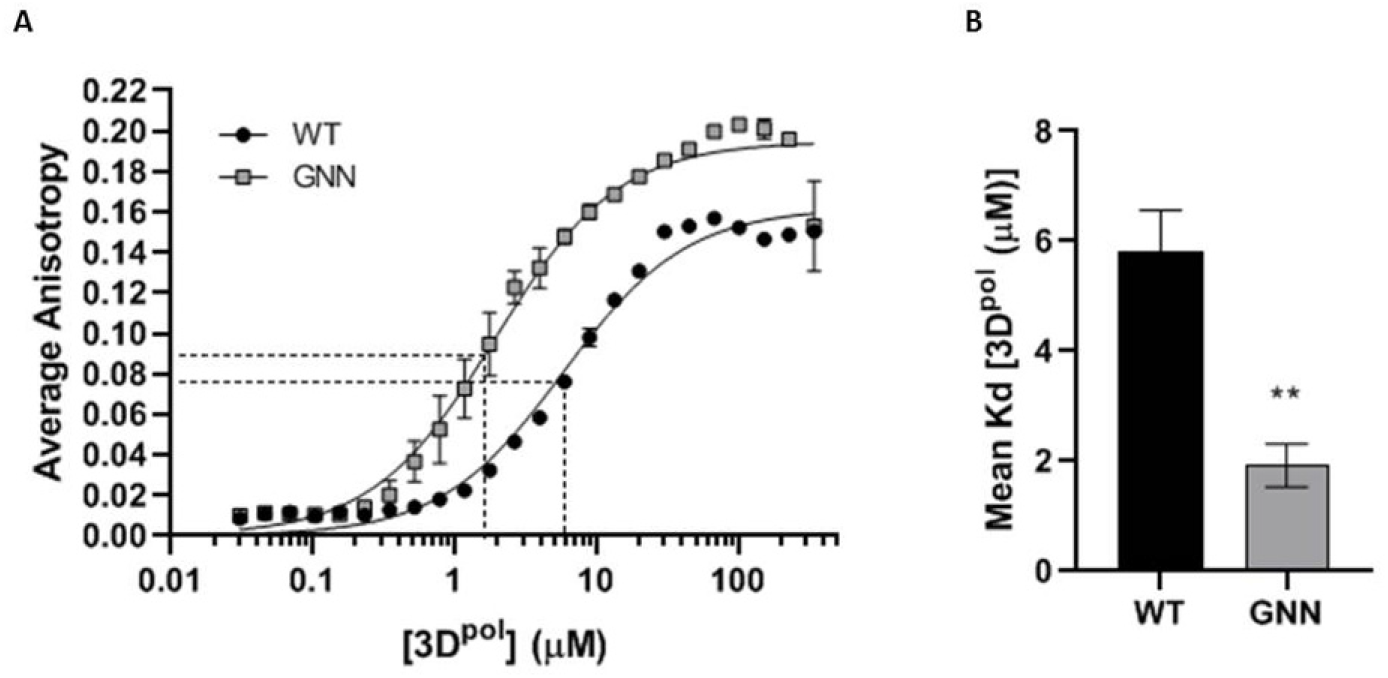
RNA binding affinity. Fluorescence polarisation anisotropy (FPA) to determine RNA binding affinities of recombinant 3D^pol^-WT and 3D^pol^-GNN A) Binding of FITC-labelled 13mer poly-A RNA to recombinant 3D^pol^ calculated as average anisotropy after 30 minute incubation. B) Mean Kd values comparing the affinity of 3D^pol^-WT and 3D^pol^-GNN with 13mer FITC-labelled poly-A RNA (n = 3 ± SEM, analysed by interpolation of a standard curve, two-tailed unpaired t-test, ** p < 0.01).

The ability of the protein to incorporate [α-32P] UTP into RNA, as described in (Bentham *et al.*, 2012) utilised a poly-A template and an oligo-d(T) primer, supplemented with a 5:1 ratio of UTP to [α-32P] UTP. After incubation at 30°C, polymerase activity was assessed by dot-blot and quantified by scintillation counting (Fig 2A, B). Normalised scintillation counts quantifying the dot-blot samples incubated for 30 minutes (Fig 2B) show a 25-fold greater incorporation of [α-32P] UTP for WT compared to the enzymatically inactive 3D^pol^-GNN.

**Figure 2.**
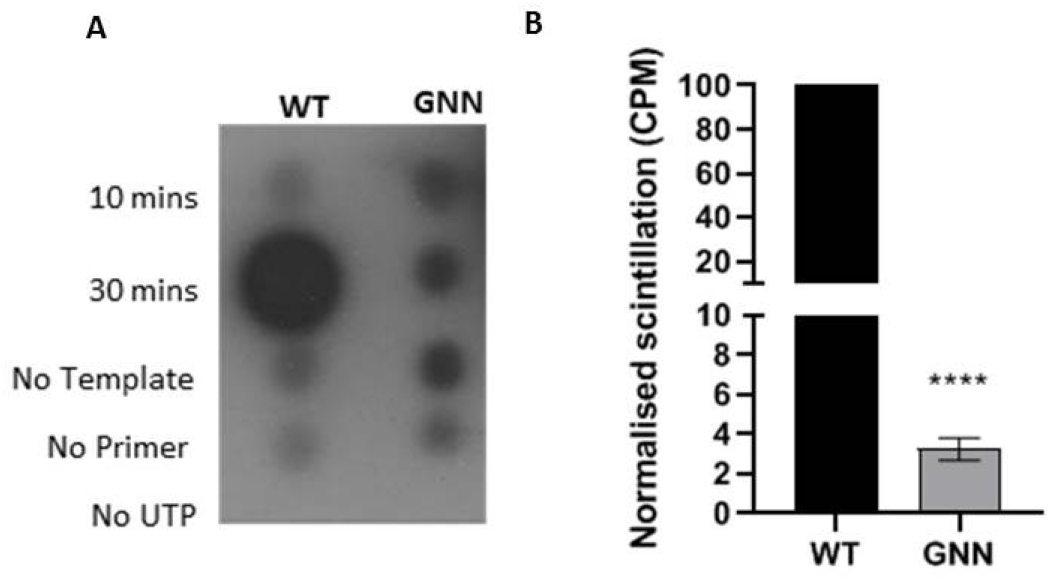
Polymerase activity assay. Polymerase activity assays were used to determine activity of recombinant 3D^pol^. A) Dot-blot showing that incorporation of [α-32P] UTP is only possible when all assay components are included. B) Quantification of levels of incorporated radiolabelled nucleotides after 30 minutes incubation by scintillation counts per minute normalised to WT (n = 3 ± SEM, analysed by two-tailed unpaired t-test, **** p < 0.0001). The GNN mutant indicates background levels.

### Formation of higher-order complexes

We have previously demonstrated the formation of higher order fibrillar complexes of 3D^pol^ by negative stain transmission electron microscopy (TEM) (Bentham *et al.*, 2012). In order to investigate the assembly of the fibrils, we employed glutaraldehyde cross-linking both to trap higher-order oligomers and stabilise the fibrils. Samples of recombinant 3D^pol^ were incubated at 30°C in the presence of glutaraldehyde, the RNA template-primer used for the activity assay described above. At time intervals, the reaction was stopped by the addition of Laemmli buffer. The samples were resolved by gradient SDS-PAGE and 3D^pol^-WT or 3D^pol^-GNN multimers detected by anti-3D^pol^ immunoblot (Fig 3A and 3C respectively) and quantified by densitometry (Fig 3B and 3D respectively). An increase in oligomer formation over time was evident, which seemed to correlate predominantly to the addition of dimers. Higher-order oligomer formation was more rapid for 3D^pol^-GNN than WT. At later timepoints, complexes were too large to migrate into the gel (Fig 3A and 3C).

**Figure 3.**
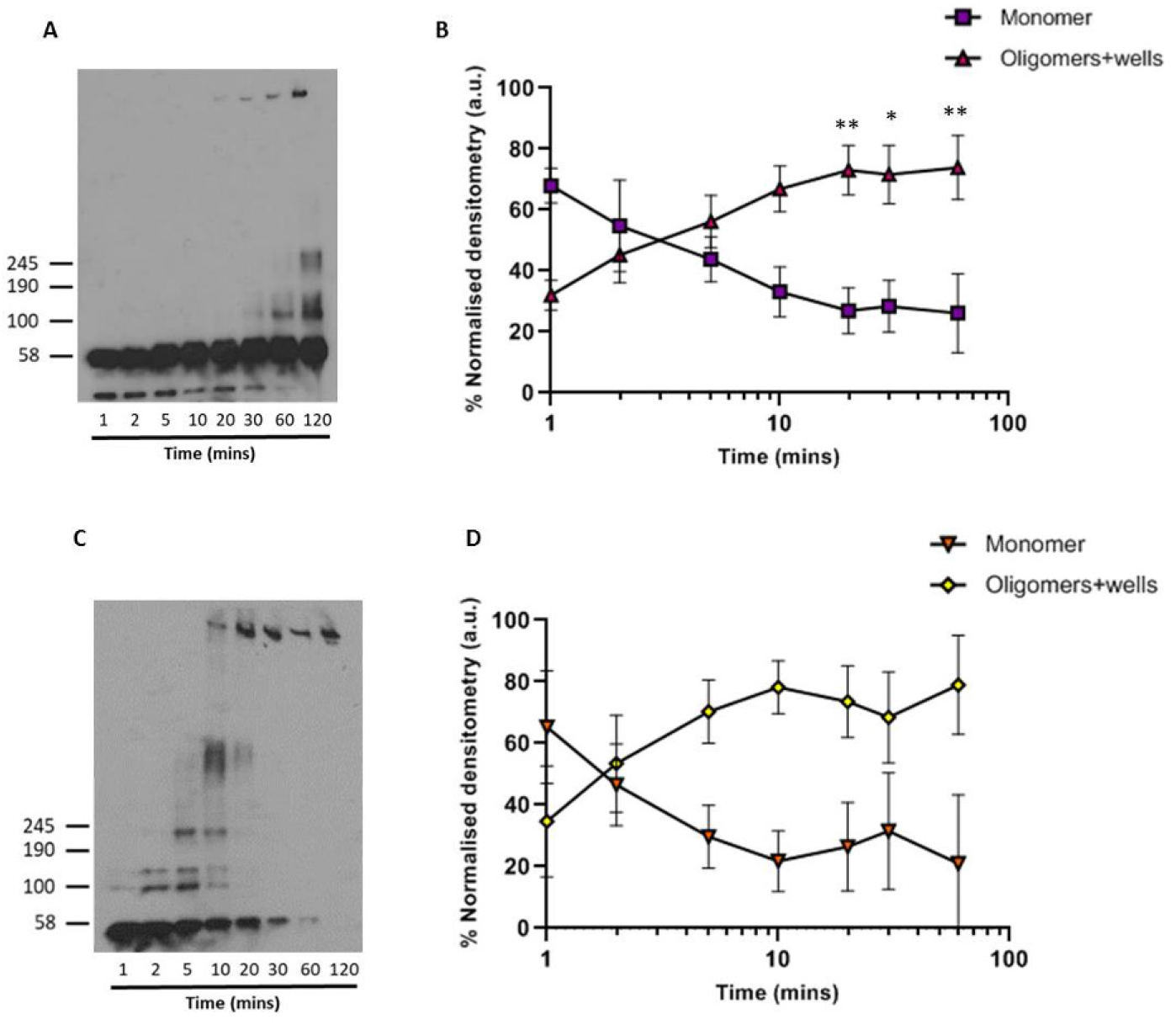
Elongation of protein monomers into higher-order oligomers. Polymerase activity assay in the presence of glutaraldehyde, showing the formation of higher-order oligomers by representative Western blot of a polymerase activity assay of A) 3D^pol^-WT and C) 3D^pol^-GNN. B) and D) Quantification of 3D^pol^ higher-order oligomer formation of 3D^pol^-WT and 3D^pol^-GNN, respectively, over time by densitometry. Values normalised to the average densitometry values (a.u) at each timepoint across three biological repeats using ImageJ. Approximate molecular weights are indicated, by comparison to Coomassie-stained gels. Comparisons between timepoints are within 3D^pol^-WT and 3D^pol^-GNN respectively (n = 3, ± SEM, analysed by two-way ANOVA, * p < 0.05, ** p <0.001).

### Structural analysis of FMDV 3D^pol^ fibrils

To investigate higher-order fibril structures in the presence of glutaraldehyde, samples of 3D^pol^-WT and 3D^pol^-GNN were taken at 30 minutes. Samples were prepared for TEM imaging using negative stain (Fig 4A and 4B). This analysis showed that 3D^pol^-WT was able to form higher-order fibrillar structures, similar to those we reported previously (Bentham *et al.*, 2012) (Fig 4A). Larger structures produced by 3D^pol^-GNN appeared as protein aggregates (Fig 4B), similar to those occasionally found in samples of 3D^pol^-WT incubated in the absence of RNA, suggesting that faster oligomerisation of 3D^pol^-GNN (shown in Fig. 3) may actually result in less ordered assembly. The availability of fixed, stable 3D^pol^ fibrils facilitated collection of high-resolution structural data to assess possible conformational heterogeneity in the population of fibrils. To this end, 3D^pol^-WT fibrils were produced as above, prepared for cryo-EM and imaged on a Thermo-Fisher Titan Krios cryo-electron microscope. Fibril sections were picked and subjected to 2D class-averaging, which appeared to show variation in fibril diameters (Fig 4C). Measurements from 1D projections of these 2D classes along the fibril axis confirmed this, showing a bi-modal distribution of diameters that ranged between ~20 nm and ~23 nm. The modal diameters were 21.3 nm and 22.6 nm, indicating the presence of at least two forms of fibril (Fig S1). These values correlate well with the size of fibrils previously measured from negative stain TEM images (Bentham *et al.*, 2012).

**Figure 4.**
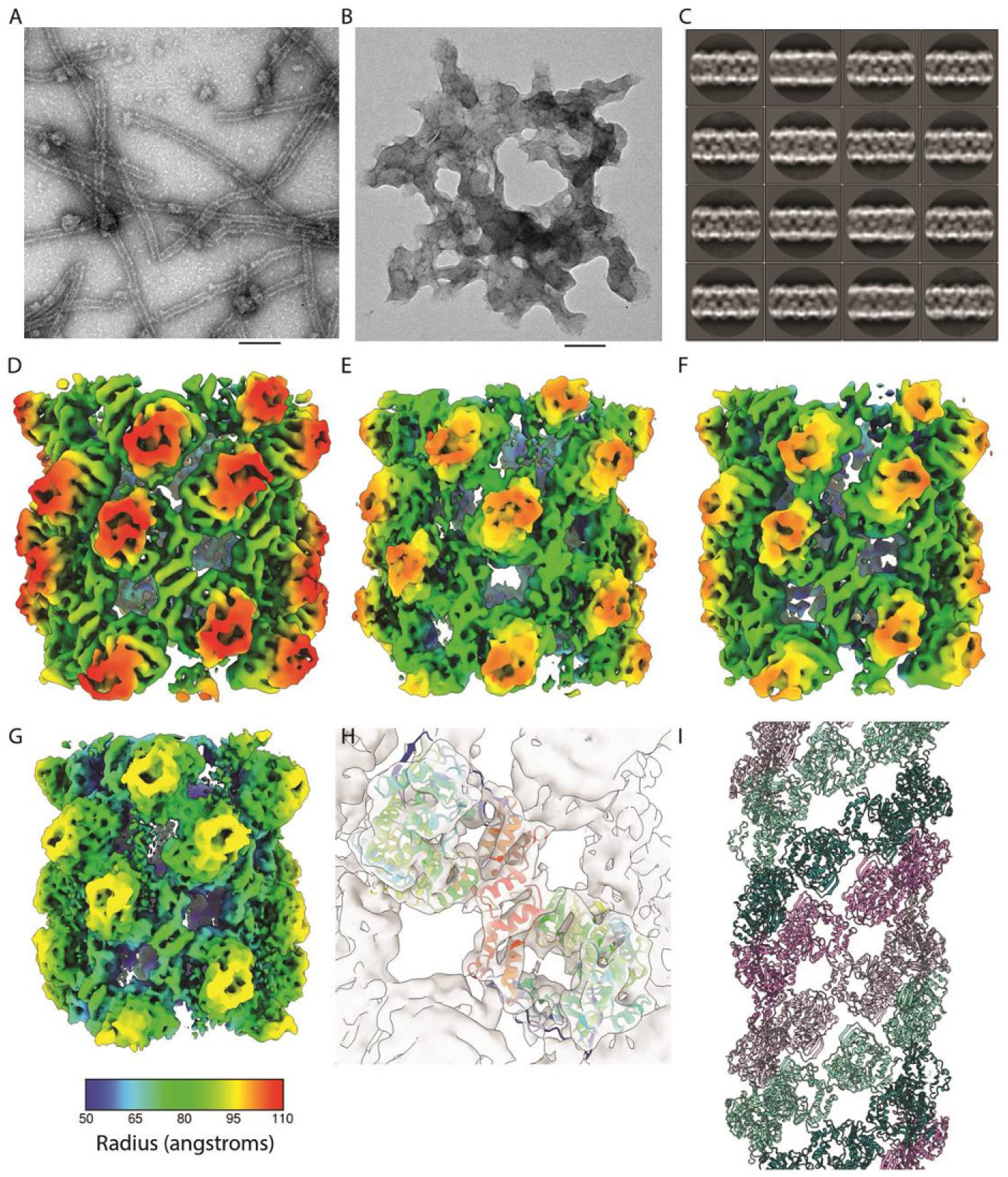
Structure of 3D^pol^ fibrils. Negative stain TEM of a sample from a polymerase activity in the presence of glutaraldehyde after 30-minute incubation. A) 3D^pol^-WT forms visible fibrils, B) 3D^pol^-GNN does not show any distinct fibril structure formation. Scale bar 100 nm. C) 2D classification and averaging of fibril section images from cryo-EM data shows variation in fibril diameter. D-G) Helical reconstruction led to the calculation of density maps for nine classes of fibril at 7.3-9.5 Å resolution, four representative maps are illustrated here coloured according to radius (see key below panel G). D) The broadest fibril reconstruction; B1 had a diameter of 22.3 nm, C2 symmetry, a helical twist of 39o and an axial rise per subunit of 27 Å. E) Conformation B2 has a more open structure, is also C2 symmetric and has a diameter of 21.7 nm, twist of 40 o and rise of 31 Å. F) N3 is a narrow (Ø = 21.5 nm) and tightly-packed fibril which may also be described by a 1-start helical symmetry with a twist of −158.8 o and axial rise of 15 Å. G) N9 is a narrow, open fibril structure having a diameter of 20 nm, twist of −158.6° and axial rise of 17.4 Å. H) Fitting of the X-ray structure for 3D^pol^-WT in complex with template primer RNA and ATP (PDB: 2EC0) into each density map shows that these fibrils are two-start helical lattices comprising protofilaments of 3D^pol^ dimers. (I) Protofilament one is coloured violet/thistle, while protofilament two is coloured teal/aquamarine, two shades are used in each protofilament to highlight how 3D^pol^ monomers form ribbons of dimers.

Helical reconstruction with 3D classification was performed, separating the dataset into fifteen self-similar classes. In this analysis, nine classes gave reconstructions that exhibited sharp density. These were individually refined leading to the definition of fibril reconstructions with diameters that reflected the bimodal distribution seen in the 2D class averages. Of these nine conformations, two were defined as ‘broad’, B1 and B2 – corresponding to the 2D classes having a modal diameter of 22.6 nm. Seven reconstructions were classed as ‘narrow’; N3-N9, corresponding to the larger group of fibrils having a modal diameter of 21.3 nm. The distinction between broad and narrow fibrils was made on the basis of the helical symmetry of each group (see below). Reconstructions ranged in resolution between 7.3 Å and 9.5 Å (Fig 4D-G, Fig S2, Table S1, Movie S1).

Each reconstruction was readily interpreted by rigid body fitting of the crystal structure for FMDV 3D^pol^ (PDB ID 2EC0) revealing that all of the solved fibril conformations are right-handed 2-start helical assemblies, each comprising a lattice of repeating 3D^pol^ dimers (Figure 4H-I). Dimers are rotationally symmetric about their interface and each of the two protofilaments is a ribbon of 3D^pol^ dimers that is therefore anti-parallel. The nine conformations that were resolved through our analysis show that variation in fibril diameter is a consequence of the two protofilaments sliding and flexing relative to each other (Movie S1). Among these reconstructions, four distinct forms of fibril were identified. The presence of a bimodal distribution of fibril diameters reflects the presence of two helical symmetries. Broad fibrils were found to be 2-start helices having C2 symmetry about the helix axis. Protofilaments had between 9 and 9.2 dimeric subunits per helix turn. Narrow fibrils were not C2 symmetric and exhibited 2-start helical symmetry with between 8.4 and 8.6 3D^pol^ dimers per helix turn in each protofilament ribbon. A notable difference between the broad and narrow classes of fibrils was that narrow fibrils could also be described by a left-handed 1-start helix of 3D^pol^ dimers having a twist of ~-159° and an axial rise of 15-17 Å while the two-fold symmetric, broad-fibrils could not (Table S1).

Within the narrow and broad classes, fibrils were found to form either tightly packed or open conformations. N3-N8 were tightly packed fibrils in which flexing of protofilaments resulted in a progressive tightening of packing from N3 to N8, with N8 being the most densely packed of the narrow fibril forms. N9 shows a rather different assembly with a longer axial rise and, although the fibril is narrower, a more open structure with large gaps between protofilaments. Similarly, B1 and B2 showed tight and open forms respectively, with B2 being narrower than B1 and having a longer axial rise per subunit. Three-dimensional classification did not distinguish more subtle variations for broad fibrils (similar to those seen in the tightly packed narrow fibril classes (N3-N8)), possibly a consequence of fewer broad fibrils present in the dataset. In both N9 and B2 (the open fibril conformations) the spaces between protofilaments contained noisy density that was not accounted for by fitting the atomic model for 3D^pol^, we postulate that this may be RNA (see below).

Supplemental movie 1 shows morphing between fibril conformations, this is intended to illustrate the differences in these structures rather than model any conformational changes that may occur. To determine whether the fibril classes solved in our analysis represented discrete structures or continuously varying helices, we traced fibril image sections from each of the reconstructions back to their original micrograph (Fig S3). We found that the narrow fibrils that contributed images to reconstructions N3-N8 varied between these forms along their length, suggesting that these reconstructions represent snapshots of a continuously flexing assembly. Fibrils were seen to switch conformation between tight and open forms. In most cases changes in fibril conformation were marked by visible constrictions and the different forms presented as discrete regions of longer fibrils, rather than the continuous variation seen for the narrow/tight classes. Overall, these data suggest that narrow and broad forms were distinct from one another. Fibrils that appear to show both broad and narrow forms along their length were, however, noted and in these images pronounced discontinuities/constrictions were observed.

### Inter- and intra-dimer interactions

As noted above, inspection of our reconstructed density maps along with rigid-body fitting of high-resolution coordinates for 3D^pol^ indicated that fibrils each comprise two protofilaments that are ribbons of dimers. Analysis of contacts and clashes between 3D^pol^ monomers in our models confirms this. These data are also consistent with fibril formation by dimer addition as shown in Fig. 3. In all conformations, the dimer interfaces and dimer-dimer interfaces are similar, while different interfaces between protofilaments were seen in tight and open forms (Fig 5A, E; Movie S2; Table S2). To describe the interactions between monomers within fibrils we use a notation incorporating the ‘palm’, ‘fingers’ and thumb’ domain nomenclature, while the relative spatial organisation of monomers within the helix are denoted as follows. Where *n* represents a monomer and then *n’* is the two-fold symmetry related monomer within a single dimeric subunit (Fig S4). Here, *n*+*1* represents the next monomer, on the opposite protofilament (and *n’*+*1* its partner within a dimer). For conformations N3-N9 the transformation from n to *n*+*1* and all subsequent increments is defined by the left-handed 1-start helical parameters: a rotation of ~-158° and an axial rise of ~15-17 Å. For the C2 symmetric broad fibrils, which do not show 1-start helical symmetry, the transformation from *n* to *n*+*1* (and subsequent odd-numbered monomers) is simply a rotation of −180°. The transformation from *n*+*1* to *n*+*2* (and all subsequent even-numbered monomers) is a rotation of ~-140° and an axial rise of ~27-31 Å. Under this notation, for all fibril conformations, *n*+*2, 4, 6* etc. are successive monomers on the same protofilament as *n*, while *n*+*1, 3, 5* etc. are on the other. The dimer interface is between thumb domains, predominantly at the C-terminal helix α16 (residues 453-467) of *n* and *n’* (Fig 5B). Dimers repeat along each protofilament ribbon with the primary interaction being between the thumb/fingertips of n and the palm domain of *n*+*2* (and between the palm domain of n’ and thumb/fingertips of *n’*+*2*; Fig 5C). Interactions between protofilaments in each conformation are also two-fold rotationally symmetric owing to the anti-parallel nature of each protofilament. The interactions are not, however, conserved between fibril forms. In tight conformations the contact interface is mainly within a loop at amino-acid residues 9-17, between *n* and *n’*+*9* or *n’*+*7* for B1 and N3-N8 respectively (Fig 5D). In conformation N8 there is an additional point of contact between *n* and *n’*+*5* between V4 in strand β1 and K65 in the N-terminal loop of the palm domain. In open fibrils the contacts/clashes between protofilaments are between *n* and *n’*+*7* for B2, and *n* and *n’*+*5* for N9. In both cases β1 strands in each monomer come together, possibly forming an extended antiparallel β-sheet (Fig. 5F).

**Figure 5.**
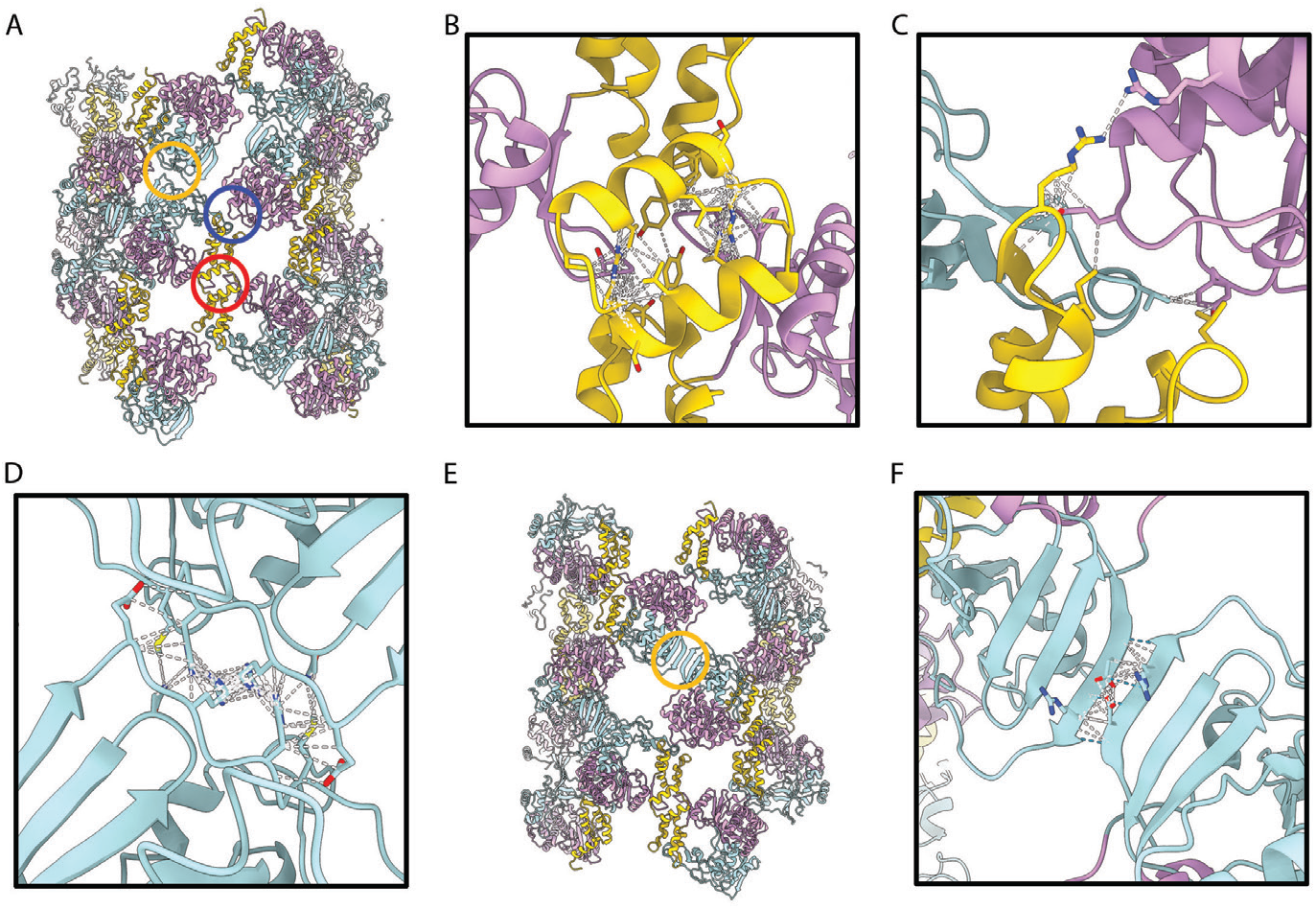
Molecular interactions within the fibril. A) The pseudoatomic model for fibril N3 is shown as cartoons coloured according to the domain structure of 3D^pol^; palm domain (plum), fingers domain (powder blue) and thumb domain (gold). Three interactions between monomers are highlighted: dimer interface (red circle), inter-dimer interface (blue circle) and inter-protofilament interface (yellow circle). Close-up views of the dimer interface (B), inter-dimer interface (C) and inter-protofilament interface are presented for C1 with contacts/clashes highlighted by white dashed lines and sidechains of putative contact residues shown. E) The pseudoatomic model of N9 is annotated to highlight the inter-protofilament interaction in this conformation (yellow circle), which is shown in close-up view with contacts/clashes highlighted with white dashed lines and putative hydrogen bonds indicated by blue dashed lines (F).

Comparison of our structures for fibrils formed of FMDV 3D^pol^ with previously published oligomeric forms of PV 3D^pol^ shows that these molecules pack differently. X-ray crystallography studies have shown that PV 3D^pol^ molecules form functional dimers via a strong interaction termed ‘interface 1’ between the C-terminal helix in the thumb domain, and the back face of the palm domain (Hansen *et al.*, 1997). Furthermore, PV helical polymerase lattices form much larger (50 nm) fibrils, helical reconstructions of which have shown a 6-start helix of protofilaments formed of interface 1 dimers (Wang *et al.*, 2013).

### Effect of 3D^pol^ mutations on replication

The cryo-EM data suggested that the fibril protofilaments are assembled via two main inter-molecular interactions, a thumb-to-thumb interaction (involving residues Y394, G395, T396, F398, P436, Y455, R456, S457, Y459, L460, V463, N464, V466, C467, D469 and A470) and a fingers/thumb-to-palm interaction (involving residues A22, P23, V25, L73, R76, H322, Y323, E324, G325, D329, T330, Y346, D347, A415, R416, R417, I420, P445, L449 and F450). It should be noted that these contact residues were identified by fitting of high-resolution data into an intermediate resolution map, thus no account is taken for potential rearrangement of side-chains upon polymerisation. Helical reconstruction of FMDV 3D^pol^ fibrils revealed considerable conformational flexibility. Many attempts were made to achieve sufficient resolution as to allow *ab initio* atomic modelling, these were not however successful.

The third interaction site (between protofilaments) is highly variable. Our fitting experiments led to the definition of contact interfaces for tight forms comprising four residues. The considerable flexing of protofilaments argues for a weak interaction at this site. The open forms show a more intriguing potential interaction – formation of an extended antiparallel β-sheet via hydrogen bonding between residues 4-8. Mutagenesis to target this interface would require disruption of secondary structure. For these reasons we have chosen to focus our mutagenesis studies on those interactions that support protofilament formation.

To understand whether the inter-molecular interactions above are important for viral replication we employed a replicon system (Tulloch *et al.*, 2014; Herod *et al.*, 2015) We hypothesised that if the polymerase inter-molecular interactions described here are biologically important, disruption of these interactions would impair function, resulting in reduced replicon replication. We therefore selected several residues from both the thumb-to-thumb interface (Y394, F398, R456, Y459, L460, V463 and C467) and the fingers/thumb-to-palm interface (P23, V25, E324, Y346, D347, I420, P445, L449) for mutation. Selecting residues at random from both interfaces prevented biasing the outcome by focusing the mutagenesis on specific secondary structures or amino acid physical properties. Each of the selected residues was individually mutated to alanine in the context of a FMDV replicon in which the structural proteins were replaced by GFP. Measuring GFP expression therefore allows the replication competency of each mutation to be assessed. From each mutant replicon, RNA was generated by T7 transcription and transfected into BHK-21 cells. Replicons with 3D^pol^-WT and 3D^pol^-GNN were included as controls, the latter for the level of translation from the input RNA. RNA replication was monitored by GFP fluorescence, with replication shown as the number of GFP positive cells at 8 hours post-transfection, analysed according to our established protocols (Herod *et al.*, 2015) (Fig 6A).

**Figure 6.**
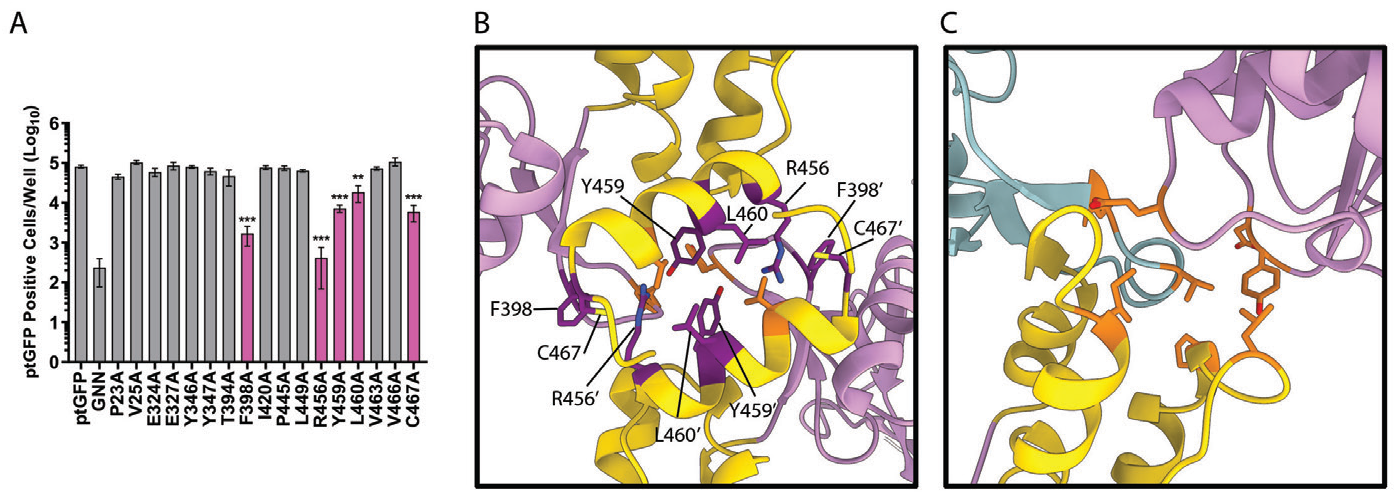
Replication of 3D^pol^ mutants. Intramolecular contact residues were identified and were mutated to alanine within the 3D-encoding region of a ptG-FP-expressing sub-genomic replicon. A) Replicon RNA was transfected into BHK-21 cells and replication determined by measuring the levels of GFP expression in comparison to the level of translation (GNN). Residues that significantly reduced the level of GFP expression are highlighted in pink. B) and C) Location of the amino-acid residues that were mutated in this analysis, those that are coloured orange did not significantly affect replicon efficiency, while those that are coloured dark magenta significantly reduced the levels of GFP expression and were located at the modelled 3D^pol^ dimerisation interface only (PDB: 2EC0). Statistical analysis by unpaired two-tailed t-test, ± SEM, ** p < 0.001, *** p <0.0001.

Of the fifteen amino acids mutated only five: F398A, R456A, Y459A, L460A and C467A, significantly reduced FMDV replicon efficiency compared to the WT replicon. All of these residues were located within the thumb-to-thumb interaction (coloured dark magenta, Fig 6B). All the other mutations tested showed levels of replication equivalent to a WT replicon (coloured orange Fig 6B and C). This included two mutations (V463A and V466A) and all of the mutations made at the fingers/thumb-to-palm interface. Overall, these data suggest that residues involved in dimer formation are important for FMDV replication, potentially through allowing formation of intermolecular 3D^pol^-3D^pol^ interactions within the FMDV replication complex.

### Unassigned density within 3D reconstructions may represent RNA

Within each of our fibril reconstructions, density within the active site of individual 3D^pol^ molecules was visible, however, this was very weak in comparison to the protein density. Nonetheless, the data suggest the presence of retained RNA at low occupancy within fibrils (Fig 7, Movie S3). In the open conformations (reconstructions B2 and N9) there is additional unassigned density that extends between two 3D^pol^ molecules, bridging an opening in the fibril between protofilaments. The density is poorly resolved but unambiguously present, we suggest that this may represent RNA entering or extruded from the active sites.

**Figure 7.**
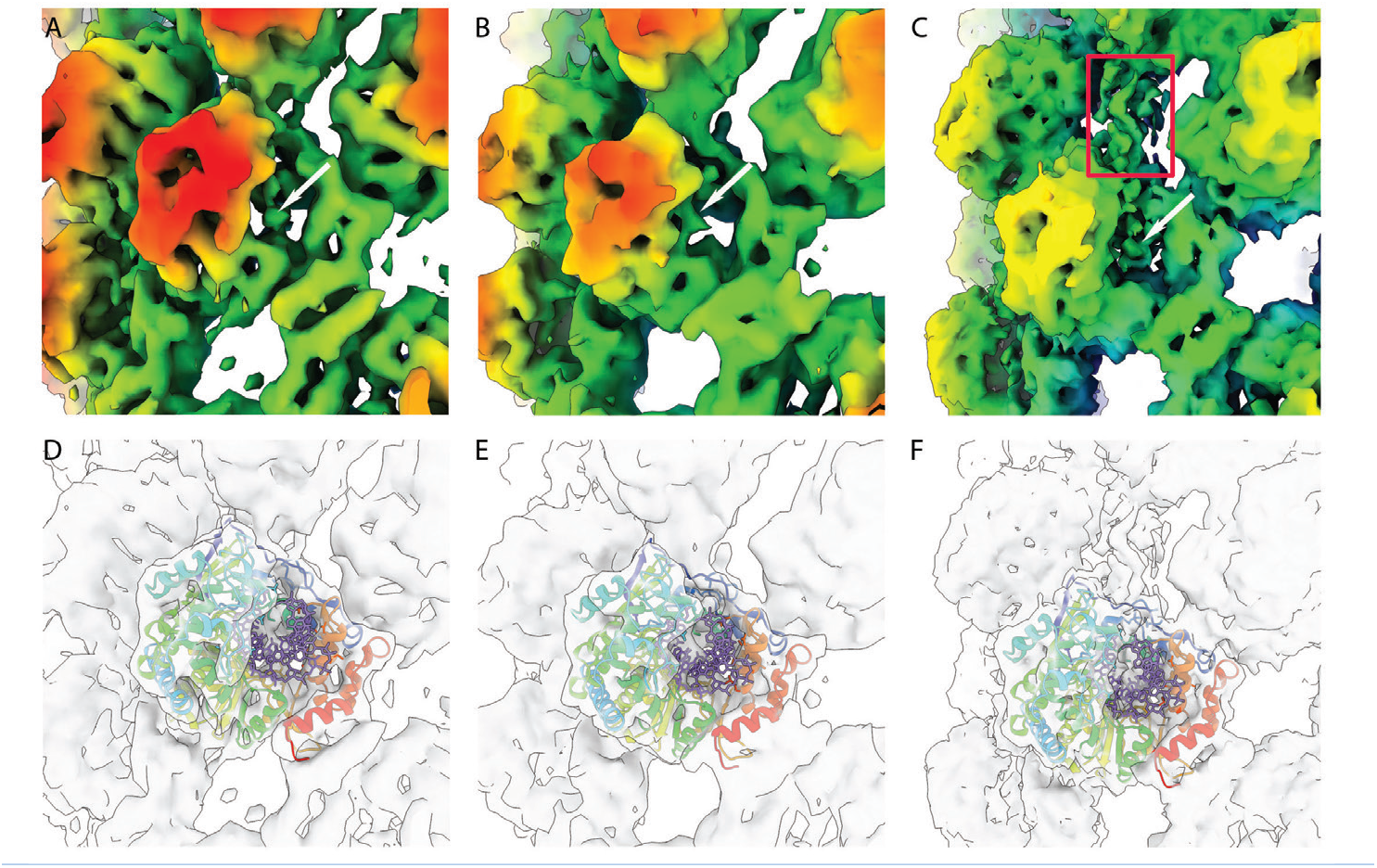
unassigned density in fibril reconstructions indicates the presence of RNA. Close up views of 3D^pol^ reconstructions for fibril B1 (A) and N3 (B) zoomed into a single monomer and coloured according to radius (key as in Fig 4F), shows the presence of weak unassigned density in the active site of 3D^pol^ that we attribute to RNA (white arrows). D, E) the same view is presented as a transparent isosurface with PDB 2EC0 shown. The atomic model is shown with protein represented as a rainbow coloured ribbon diagram and RNA as sticks coloured mauve, to highlight the assignment of protein and RNA to cryo-EM density. C, F) Conformation N9 also shows unassigned density in the active site as well as a finger of density linking two protofilaments (red rectangle) that may be RNA template or product extruded from the active site.

## Discussion

The RNA-dependent RNA polymerase, 3D^pol^, was one of the first proteins to be characterised both in PV (Baltimore and Franklin, 1962) and in FMDV (Polatnick and Arlinghaus, 1967) and is the key enzyme in viral replication. Both PV and FMDV 3D^pol^ have been shown to form higher-order structures in vitro in the form of fibrils (Pata *et al.*, 1995; Lyle *et al.*, 2002; Spagnolo *et al.*, 2010; Tellez *et al.*, 2011; Bentham *et al.*, 2012; Wang *et al.*, 2013) that have been postulated to be part of viral replication complexes. It would be expected that replication complexes would be dynamic structures, and although there is structural data available for the PV fibrils this is at relatively low resolution, did not include any RNA and was thus unable to give insight into any dynamics (Tellez *et al.*, 2011; Wang *et al.*, 2013). Here, we report multiple conformations of the fibrils formed by FMDV 3D^pol^, some of which occur on the same molecule, and evidence for RNA within the complex. We probed the interfaces by mutagenesis and propose a model for the potential role of dynamic changes in the fibril during genome replication.

After demonstrating that the recombinant 3D^pol^ protein used here was able to bind RNA and was catalytically active, we analysed the structures of assembled products of a polymerase activity assay after crosslinking. This provided a total of 9 different 3D^pol^ reconstructions, all of which are at higher resolution and distinct from the previously reported structures of PV fibrils, in terms of both dimensions and the orientation of molecules within the helix. The dimerisation interface of the FMDV 3D^pol^ molecules is novel and distinct from the dimerisation of PV 3D^pol^ molecules. Formation of fibrils by dimer addition is also supported by the cross-linking data presented here. Mutations that affected the ability of the polymerase to transcribe replicon RNA were introduced within the dimerisation thumb-thumb interface. This contact region has been implicated in the formation of the PV fibrils and shown to be necessary in the production of virus, despite being distant from the active site (Watkins *et al.*, 2020). In PV, a model has been described whereby RNA would travel through the active site between polymerase molecules, however, RNA was not detected (Lyle *et al.*, 2002; Tellez *et al.*, 2011). In support of this model, density corresponding to RNA was detected in the FMDV fibril structures presented here. This density appears in the active site of all our reconstructions and is also seen linking individual molecules in the open fibril conformations.

The formation of a classical intracellular, membrane-associated replication complex for FMDV has yet to be demonstrated within infected cells. One could speculate, in the context of a cellular replication complex, that the polymerase molecules could coat newly synthesised negative-sense intermediates and prevent the formation of double-stranded RNA, a potent trigger of an innate immune response. Avoiding double-stranded RNA formation would also allow for a more energetically-favourable replication process. The addition of dimers to coat RNA strands could generate helical fibrils similar to those described here. The concept of 3D^pol^ acting in a similar way to the N protein of negative sense viruses or the coat protein of helical viruses such as tobacco mosaic virus is intriguing.

The structures described here could alternatively represent trapped conformers of the active replication machinery within the functional complex. The different fibril conformations observed could represent different stages of replication. Although there is some density for RNA in the active site of 3D^pol^ molecules within the fibrils, a model where RNA is looping out from the body of a dynamic fibril would be consistent with the relatively weak RNA density seen overall. However, it is also tempting to speculate that the density seen between 3D^pol^ molecules in the open N9 conformation correlates with newly synthesised RNA being extruded from the active site.

The 9 fibril structures described here are broadly representative of 4 types: narrow, broad, tight and open. Data for the narrow fibrils were consistent with these being a continuous, flexing assembly, with the broad fibrils a separate, distinct class. Given the breadth of fibril morphology shown here, it is therefore possible that both models described above are correct. Some of the fibrils could form by the simple addition and dissociation of 3D^pol^ subunits and act to coat newly formed RNA transcripts. Others could form part of a dynamic replication complex. Different pools of 3D^pol^ molecules involved in replication would be consistent with the models we proposed in (Herod et al, 2016) as a result of complementation experiments. Correlative light-electron microscopy studies in replicon-containing cells are underway in order to further elucidate the formation and dynamics of complexes within cells and to establish if this and other non-structural proteins, such as 3CD, 3A and 3B, are able to interact with the FMDV 3D^pol^ fibrils.

## Materials and Methods

### Expression and purification of his-tagged 3D^pol^

WT C-terminally His-tagged 3D^pol^ was expressed from vector pET-28a (a kind gift from Esteban Domingo, Centro De Biologia Molecular Severo Ochoa, Madrid, Spain) (Arias *et al.*, 2005) in Codon plus (+)BL21 using methods described previously (Ferrer-Orta *et al.*, 2004). For lysis, the cells were suspended in 20 mM Tris-HCl [pH 8] lysis buffer with 20 mg/ml lysozyme before sonication as described previously (Ferrer-Orta *et al.*, 2004). Lysates were clarified and the supernatant filtered through 0.22 μm filters and subjected to nickel affinity chromatography using 1 ml HisTrap HP columns (GE Healthcare). Elution was carried out via a 25-500 mM imidazole gradient in 50 mM Tris [pH 8], 500 mM NaCl buffer. Fractions containing the protein were pooled, dialyzed and stored in aliquots at −20 °C as described previously (Herod *et al.*, 2016). A replication-deficient enzyme (termed GNN which includes the active site mutations D338N and D339N (Herod *et al.*, 2016) was expressed and purified in the same way and the mutations confirmed by mass spectrometry.

### 3D^pol^ RNA-binding by fluorescence polarisation anisotropy

The ability of FMDV 3D^pol^ to bind to RNA was examined using fluorescence polarisation anisotropy (FPA) adapted from published protocols (Selvaraj *et al.*, 2018). FPA measures the tumbling rate of a fluorescently labelled molecule. After excitation by polarised light, the excited fluorophore emits light with the degree of polarisation inversely proportional to the tumbling rate of the fluorescent molecule in solution. Therefore, the smaller the molecule, the higher the tumbling rate resulting in the depolarisation of the emitted light as the fluorophore reorients in solution during the lifetime of the excitation. FPA was undertaken in a black opaque 384-well format (Perkin Elmer). 20 μl of RNA binding buffer (20 mM Tris-HCl (pH 8), 100 mM NaCl, 0.01 % Triton X-100) was added to each well followed by a serial dilution of purified 3D^pol^ or mutant 3D^pol^-GNN. 20 μl of 20 nM 3’FITC-labelled 13mer poly-A (Dharmacon) (kind gift from Thomas Edwards, University of Leeds) was overlaid into each well and samples incubated at room temperature for 1 hour. Control samples that contained no fluorescent RNA to establish background polarisation were included. Polarisation was measured using a Tecan plate reader with a 480-nm excitation filter and 530 nm S and P channel emission filters.

### 3D^pol^ Activity Assay

Activity of purified FMDV 3D^pol^ was assessed by the ability of the protein to incorporate [α-32P] UTP into nascent RNA transcripts, as previously described (Ellingham *et al.*, 2006; Bentham *et al.*, 2012). The final concentration of 3D^pol^ that was included in the assays was 18.52 μM. Reactions were stopped by the addition of 1.5 μl 500 mM EDTA at the indicated times. For scintillation, 2 μl of each reaction was blotted onto filter paper (Whatmann) and allowed to dry prior to being washed as described previously (Ellingham *et al.*, 2006). The dry filter paper was exposed to film for imaging before the individual dots on the filter paper were excised and scintillated with the addition of 3 ml scintillation fluid (Perkin Elmer). Radioactive counts were measured as scintillation counts per minute.

For glutaraldehyde crosslinking, 3D^pol^ activity assays, substituting [α-32P] UTP with 4 mM UTP were employed. Samples were incubated with 5mM glutaraldehyde at 30°C for increasing time periods and reactions stopped by the addition of 30 μL 2x Laemmli buffer. The samples were denatured and analysed by gradient SDS-PAGE (12 −4 % resolving gels, and 4 % stacking gel). Immunoblotting was performed as described previously (Forrest *et al.*, 2014) with primary rabbit anti-3D 397 polyclonal antibody (kind gift from Francisco Sobrino, Centro De Biologia Molecular Severo Ochoa, Madrid, Spain) and detected with anti-rabbit HRP (Sigma-Aldrich).

### Transmission electron microscopy (TEM)

Carbon-coated copper grids were prepared by glow discharge at 25 mA for 30 seconds. The activity assay master mix was prepared as described for the crosslinking assay. Samples were incubated for 30 minutes at 30°C and then immediately 10 μl was adsorbed onto carbon-coated copper EM grids for 1 minute. Excess sample was removed and the grids washed twice with 10 μl dH2O for 5 seconds prior to staining with 10 μl 2% uranyl acetate for 1 minute, the excess was then removed with filter paper. TEM was performed on a JEOL 1400 (Jeol USA Inc.) at 30,000 x magnification.

### Cryo-electron microscopy (Cryo-EM)

Activity assay master-mix was prepared as described above and incubated at 30°C for 30 minutes. 10 μl of the master-mix was loaded on a C-flat (protochips) CF1.2/1.3-4C grid (1.2 μm holes) that had been prepared by addition of a continuous carbon support film and glow discharged at 25 mA for 1 minute. The grids were blotted for 6 seconds at blot force 10, using an FEI Vitrobot Mark IV in chamber conditions of 4°C and 95 % relative humidity. These were screened and initial analysis performed using a JEOL 2200 TEM equipped with a Direct Electron DE20 camera. Optimised grids were then imaged on a Thermo-Fisher Titan Krios in the Astbury Biostructure Laboratory using a Falcon III detector in integrating mode operated at 40 frames per second. Data were collected at nominal 75,000 x magnification, giving a sampling frequency of 1.065 Å/pixel. Two second exposures were collected at a dose rate of 54 e^−^/Å2 /second and a defocus range of −1.5 to 2 microns.

### Image processing

Preliminary analysis of a small dataset collected on a JEOL 2200 was performed using Relion 2.0.3. A single 2D class average was used to generate a starting model using SPIDER. Initial estimates of approximate helical parameters were derived by taking measurements from 2D class averages (twist = 40° and rise per subunit of 29 Å). Calculation of an initial helical reconstruction was performed giving rise to an interpretable density map with twist = 42.4° and rise of 30 Å. Inspection of this preliminary map revealed a two-start helical assembly (Fig. S5).

Collection of the substantially larger dataset was undertaken at the Astbury BioStructure Laboratory. 5417 micrographs were recorded from which 266,263 images of fibril sections were selected. Motion correction was performed using Motioncor2 with dose-weighting (Zheng *et al.*, 2017). CTF estimation was performed using GCTF (Zhang, 2016). Helical reconstruction was performed using Relion 3.0 followed by 3.1 (Scheres, 2012; He and Scheres, 2017). Following 2D classification the dataset was reduced to 247,623 fibril section images that had contributed to sharp class averages. These 2D class average images were carefully realigned to ensure that in all images the helix was parallel to the Y axis, using SPIDER (Frank *et al.*, 1996), they were then projected to one dimension (1D) using Bsoft (Heymann, 2001). Radial density for each class average was plotted using Prism 8 (GraphPad), and fibril diameters were estimated as the distance between the minima on each side of the helix axis. Based on this 2D classification analysis, a dataset of fibril images that gave sharp class averages was selected. 3D classification was performed with k = 15, leading to the definition of nine classes of fibrils having diameters that reflected the full distribution of diameters identified by measurements of the radial density profiles. Inspection of the resulting reconstructions revealed that maps corresponding to the narrower class of fibril (Ø = 20-21.5 nm) could be described as left-handed 1-start helices with twist values of ~-158.8° and axial rise values of ~15-17 Å. The broader class of fibrils (Ø = 21.7-22.3nm) were two-fold symmetric about the helix axis and were processed as right-handed 2-start helices (twist= ~40°, axial rise = 27-31 Å) with C2 symmetry imposed. 3D automated refinement led to calculation of density maps for each class at resolutions ranging between 7.6 and 10.9 Å. As the reconstructions comprised an asymmetric unit that was a dimer, there was a two-fold symmetry present that could not be accommodated within the helical reconstruction process. This was subsequently imposed on half-maps using UCSF Chimera (Pettersen *et al.*, 2004). Briefly, two copies of each half-map were opened, one was rotated 180° about the x axis (perpendicular to the helix axis), the fit in map function was then used to align the two maps and the density was averaged. Symmetrized half-maps were then imported into Relion for post-processing analysis, resolution estimates ranged between 7.3 and 9.5 Å.

Fibril images contributing to each reconstruction were traced back to their original micrographs using rln-parts-trace.py (https://github.com/attamatti/relion_particle_backtrace).

Reconstructed density maps were interpreted by docking of published coordinates for the FMDV 3D^pol^ protein using UCSF Chimera (Pettersen *et al.*, 2004) and visualised using UCSF ChimeraX (Goddard *et al.*, 2018)

### Replicon plasmids

The FMDV replicon plasmid, pRep-ptGFP and the replication-defective 3D^pol^ GNN mutant derivative, have been described previously (Herod *et al.*, 2015). In this replicon, the capsid protein-coding region has been replaced with a green fluorescent reporter protein. Mutations were introduced into the pRep-ptGFP replicon 3D^pol^ coding region by overlapping PCR site-directed mutagenesis. The sequences of primers are available on request.

### FMDV replicon replication assays

To generate in vitro transcribed RNA for transfection, replicon plasmid DNA linearized with AscI (NEB, Ipswich, Massachusetts, USA) was used to template T7 in vitro transcription reactions, as previously described (Herod *et al.*, 2015). T7 RNA transcripts were purified using a Zymogen RNA Clean and Concentrator-25 (Zymo Research, Irvine, California USA) according to manufacturer’s instructions and the RNA quality was determined by MOPS-formaldehyde agarose gel electrophoresis.

To assay replicon replication, BHK-21 cells seeded into 24-well tissue cultures plates were allowed to adhere for 16 hours before transfection in duplicate with in vitro transcribed replicon transcripts using Lipofectin reagent as previously described (Herod *et al.*, 2015). Fluorescent reporter protein expression was monitored using an IncuCyte Zoom Dual Colour FLR (Essen BioSciences), within a 37°C humidified incubator, scanning hourly up to 24 hours post transfection. Images were captured and analysed using the associated software for fluorescent protein expression. Control transfections (untransfected and the 3D^pol^-GNN transfection to assess the level of input translation) were used to determine fluorescent thresholds and identify positive objects from background fluorescence. The number of positive cells per well was determined from the average of up to nine non-overlapping images per well.

## Acknowledgements

We would like to thank Professor Francisco Sobrino for the generous gift of antibodies, Professor Esteban Domingo for the generous gift of the recombinant protein, and Dr Thomas Edwards for reagents. We thank Danielle Pierce and Veronica DeJesus for technical assistance with mutagenesis and Professors Dave Rowlands and Neil Ranson for helpful comments on the manuscript.

E.A.L Investigation, data analysis, methodology, writing original draft

D.B. Investigation, data analysis, funding acquisition, data curation, methodology, supervision, visualisation, writing original draft.

J.S. Investigation, data analysis.

M.H. Investigation, data analysis, funding acquisition, supervision, editing manuscript

M.R.H. Investigation, supervision, editing manuscript

R.T. Investigation, data analysis, editing manuscript

N.J.S. Investigation, funding acquisition, supervision, writing original draft.

CryoEM maps have been deposited in the EM-dataank (https://www.ebi.ac.uk/pdbe/emdb/) with the accession numbers (B1) EMD-12093, (B2) EMD-12094, (N3) EMD-12096, (N4) EMD-12097, (N5) EMD-12098, (N6) EMD-12099, (N7) EMD-12100, (N8) EMD-12101, (N9) EMD-12102. The raw micrograph movies are deposited in the EMPIAR databank with accession number 10602.

