## Supplemental table 1 for "Higher-order structures of the foot-and-mouth disease virus RNA-dependent RNA polymerase required for dynamic inter-molecular interactions involved in viral genome replication"

**Imaging and reconstruction parameters**

| Reconstruction | B1 | B2 | N3 | N4 | N5 | N6 | N7 | N8 | N9 |
| --- | --- | --- | --- | --- | --- | --- | --- | --- | --- |
| Microscope | Thermo Fisher Scientific Titan Krios | | | | | | | | |
| Detector | Thermo Fisher Scientific Falcon III (linear mode) | | | | | | | | |
| Pixel size in micrographs | 1.065 Å | | | | | | | | |
| Pixel size in reconstruction | 2.13 Å | | | | | | | | |
| Electron fluence | 54 e/Å^2^/s | | | | | | | | |
| Frames/Movie | 79 | | | | | | | | |
| Exposure time | 2s | | | | | | | | |
| Number of Micrographs | 5417 | | | | | | | | |
| Particles picked | 266,263 | | | | | | | | |
| Particles used | 38,886 | 6,903 | 15,167 | 26,032 | 10,714 | 14,427 | 29,141 | 18,808 | 16,461 |
| Fibril diameter | 22.3 nm | 21.7 nm | 21.5 nm | 21.3 nm | 20.9 nm | 20.9 nm | 20.7 nm | 20.7 nm | 20 nm |
| Symmetry | C2 | C2 | C1 | C1 | C1 | C1 | C1 | C1 | C1 |
| Helical twist | 39.003° | 40.0259° | -158.874° | -159.002° | -158.879° | -158.778° | -158.811° | -158.545° | -158.581° |
| Rise per subunit | 26.9991 Å | 31.2392 Å | 15.0274 Å | 15.0837 Å | 15.0192 Å | 14.9352 Å | 15.1039 Å | 14.948 Å | 17.3913 Å |
| Map resolution | 7.3 Å | 9.5 Å | 8.5 Å | 7.7 Å | 7.6 Å | 7.6 Å | 7.3 Å | 7.5 Å | 7.3 Å |
