## Supplemental table 2 for "Higher-order structures of the foot-and-mouth disease virus RNA-dependent RNA polymerase required for dynamic inter-molecular interactions involved in viral genome replication"

A. Contact residues at the dimer interface

| Conformation 1 *n* | Conformation 1 *n’* |
| --- | --- |
| **Y394, G395, T396**, F398, P436, **Y455, R456, S457, Y459, L460, V463**, **C467, D469, A470** | **Y394, G395, T396**, F398, P436, **Y455, R456, S457**, **Y459, L460, V463**, **C467, D469, A470** |
| Conformation 2 *n* | Conformation 2 *n’* |
| **Y394, G395, T396**, F398, P436, **Y455, R456, S457**, **Y459, L460, V463**, V466V, **C467, D469, A470** | **Y394, G395, T396**, G397, F398, P436, **Y455, R456, S457**, **Y459, L460, V463**, V466, **C467, D469, A470** |
| Conformation 3 *n* | Conformation 3 *n’* |
| **Y394, G395, T396**, F398, **Y455, R456, S457**, **Y459, L460, V463**, **C467, D469, A470** | **Y394, G395, T396**, **Y455, R456, S457**, **Y459, L460, V463**, **C467, D469, A470** |

B. Contact residues at the dimer-dimer interface

| Conformation 1 *n* | Conformation 1 *n+2* |
| --- | --- |
| **A22, V25, A415, R416, R417, I420, P445, L449** | **R76, E324, G325, Y346** |
| Conformation 2 *n* | Conformation 2 *n+2* |
| **A22,** P23, **V25**, G28, **A415, R416, R417, I420, P445, L449,** F450 | L73, **R76,** H322, Y323, **E324, G325,** D329, T330, **Y346,** D347 |
| Conformation 3 *n* | Conformation 3 *n+2* |
| **A22, V25, A415, R416, R417, I420, P445, L449,** F450 | **R76, E324, G325,** T330**, Y346** |

C. Contact residues at the protofilament interface

| Conformation 1 *n* | Conformation 1 *n’+5* |
| --- | --- |
| E10, R12, H14, M16, N287 | E10, R12, H14, M16, N287 |
| Conformation 2 *n* | Conformation 2 *n’+3* |
| V4, K65 | V4, K65 |
| Conformation 2 *n* | Conformation 2 *n’+5* |
| R12, H14, M16, N287 | R12, H14, M16, N287 |
| Conformation 3 *n* | Conformation 3 *n’+3* |
| I3, V4, D5, T6, R7, D8 | I3, V4, D5, T6, R7, D8 |
