## Supplemental figures for "Higher-order structures of the foot-and-mouth disease virus RNA-dependent RNA polymerase required for dynamic inter-molecular interactions involved in viral genome replication"

**Supplemental data**

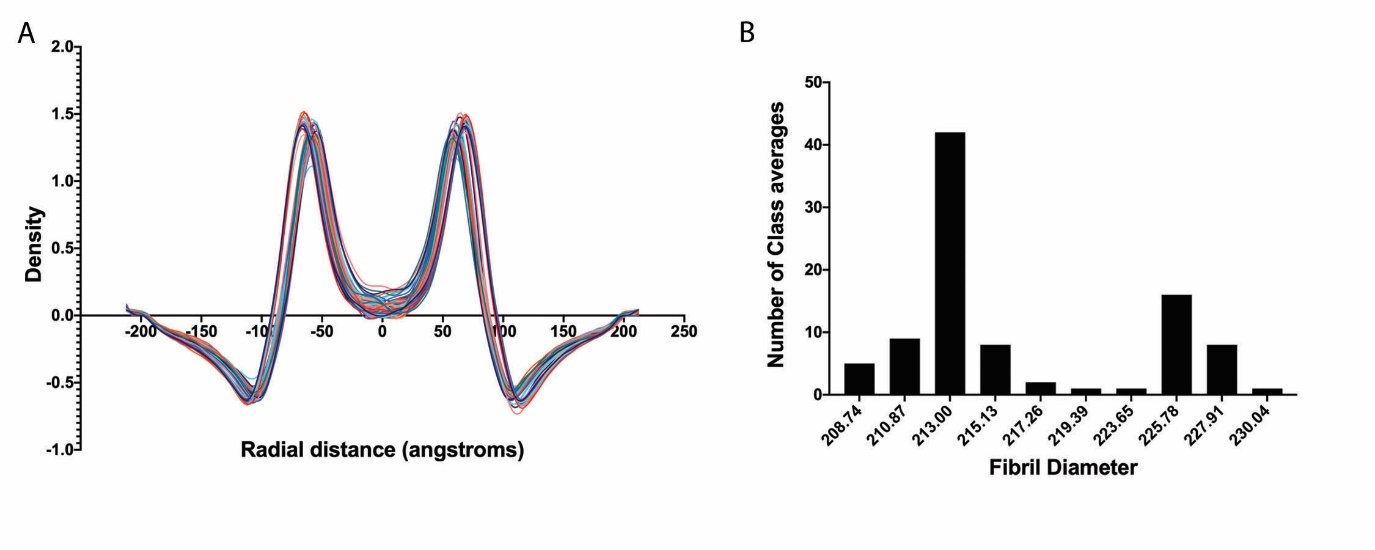

**Figure S1**

A) Radial density profiles for 100 2D class averages calculated from ~250,000 fibril image sections. Each class average was projected along the helix axis, summing the density into a 1D profile. For ease of interpretation x-axis values are given as distances from the centreline of the 2D class average images, as an approximation of the helix axis position. B) To estimate the fibril diameters for each class average image, the distance between minima in each radial density plot was calculated, a histogram of these data shows a bimodal distribution of fibril diameters ranging between 20.8 and 23 nm.

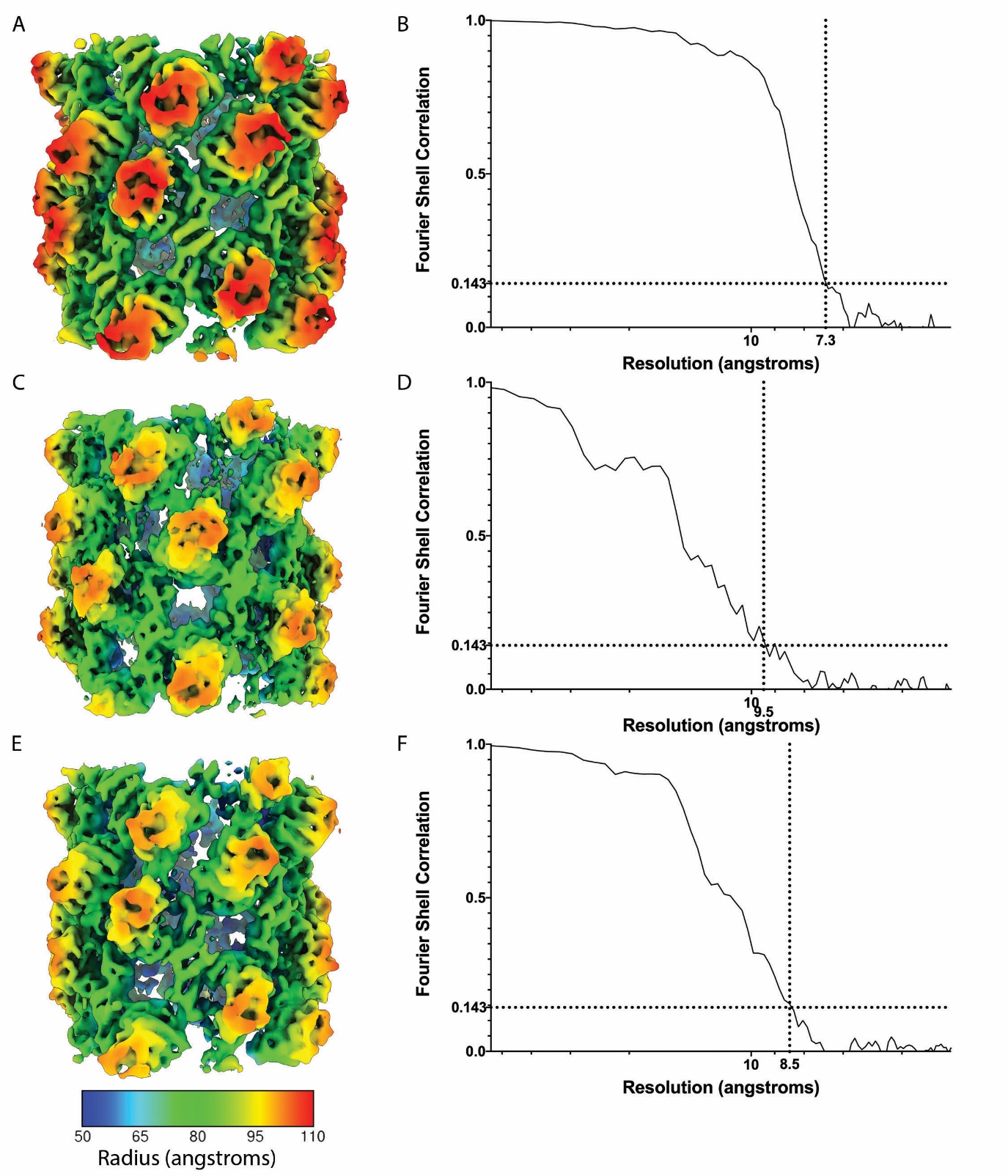

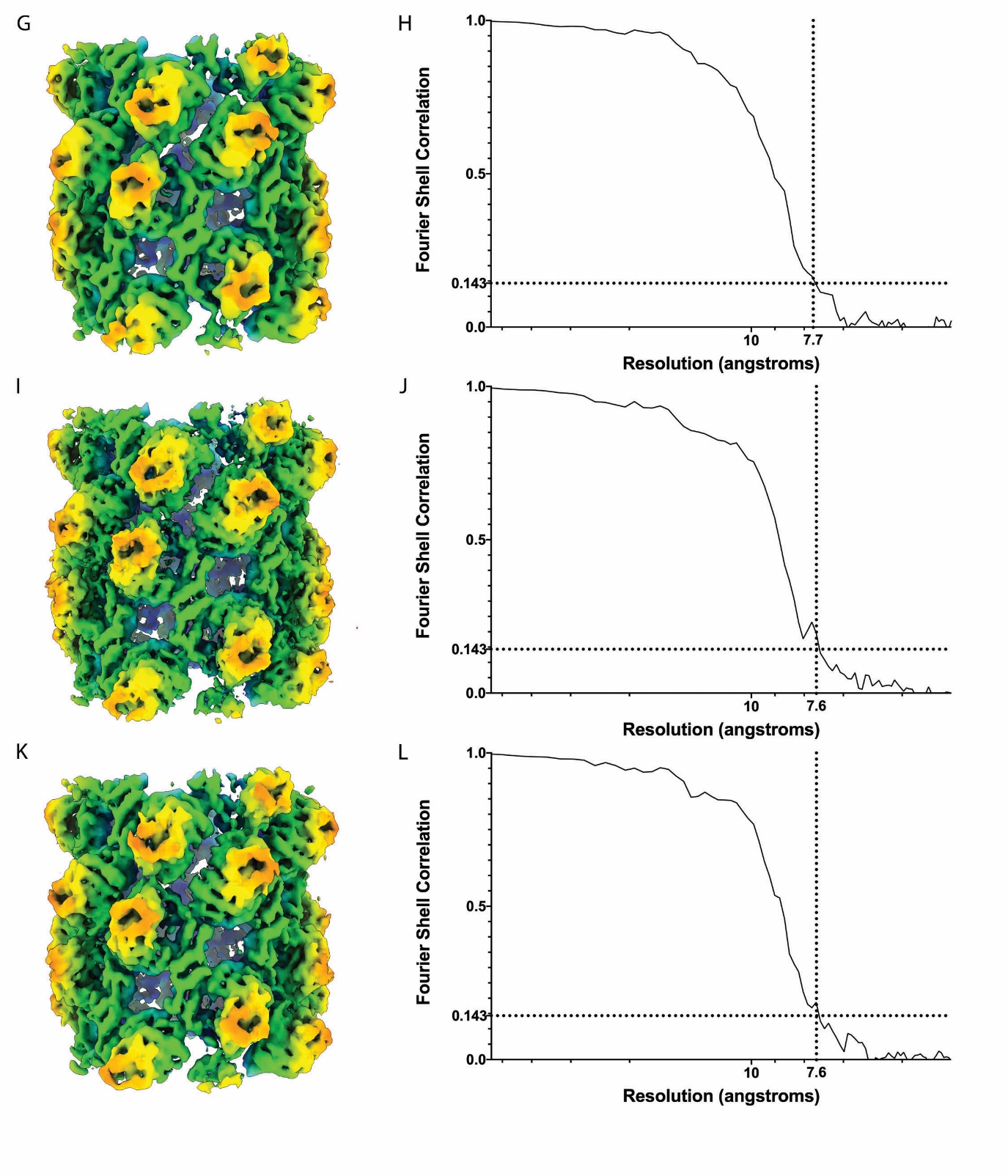

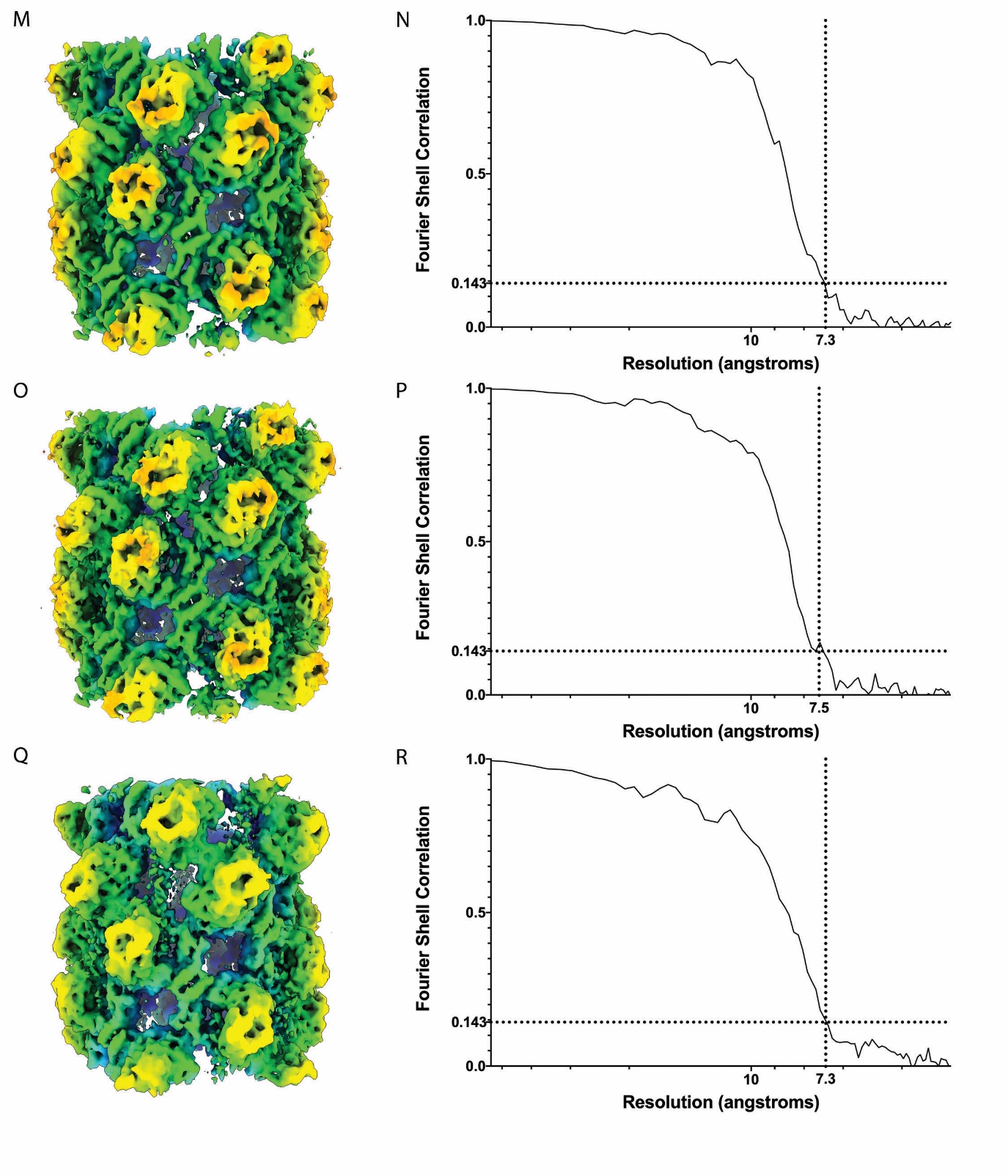

**Figure S2**

Radial coloured 3D isosurface representations and Fourier shell correlation plots for the each cryo-EM 3D reconstruction. Resolution was estimated as the point at which correlation between two half-maps, calculated independently using Relion according to the ‘gold-standard’ method, drops below 0.143. (A, B) B1, (C, D) B2, (E, F) N3, (G, H) N4, (I, J) N5, (K, L) N6, (M, N) N7, (O, P) N8, (Q, R) N9.

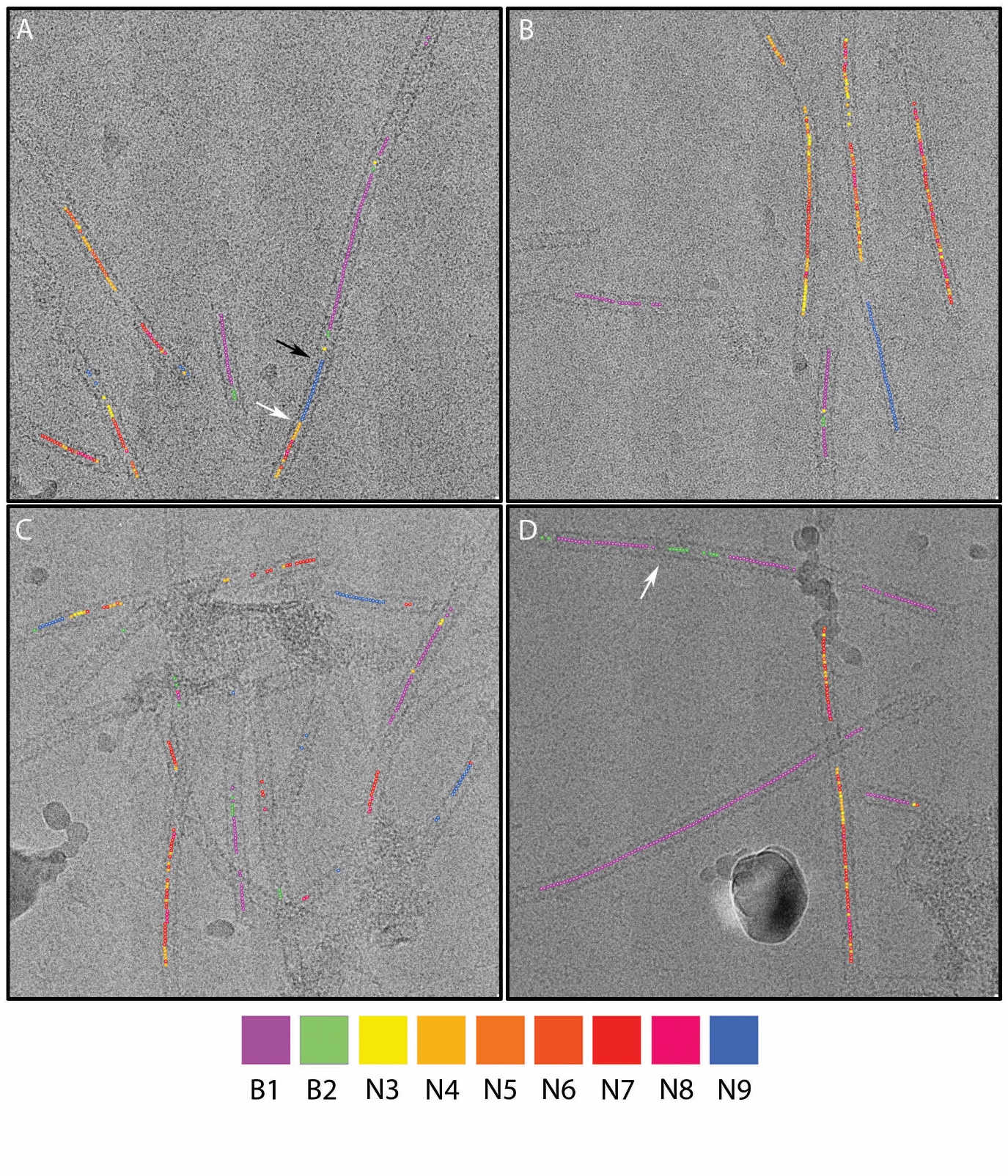

**Figure S3**

Four micrographs are shown annotated to show the position of fibril image sections that contributed to each of the three reconstructions. This shows that fibrils may comprise regions in all three states, however, while narrow/tight-packed forms are seen to alternate along single fibrils, open forms are seen in discrete sections of longer fibrils. In general broad and narrow conformations appear to be distinct, some instances of apparent transition from one narrow to broad are seen, but are accompanied by discontinuities in the density (A-black arrow), changes from tight to open forms are marked by the presence of constrictions (A,D – white arrows).

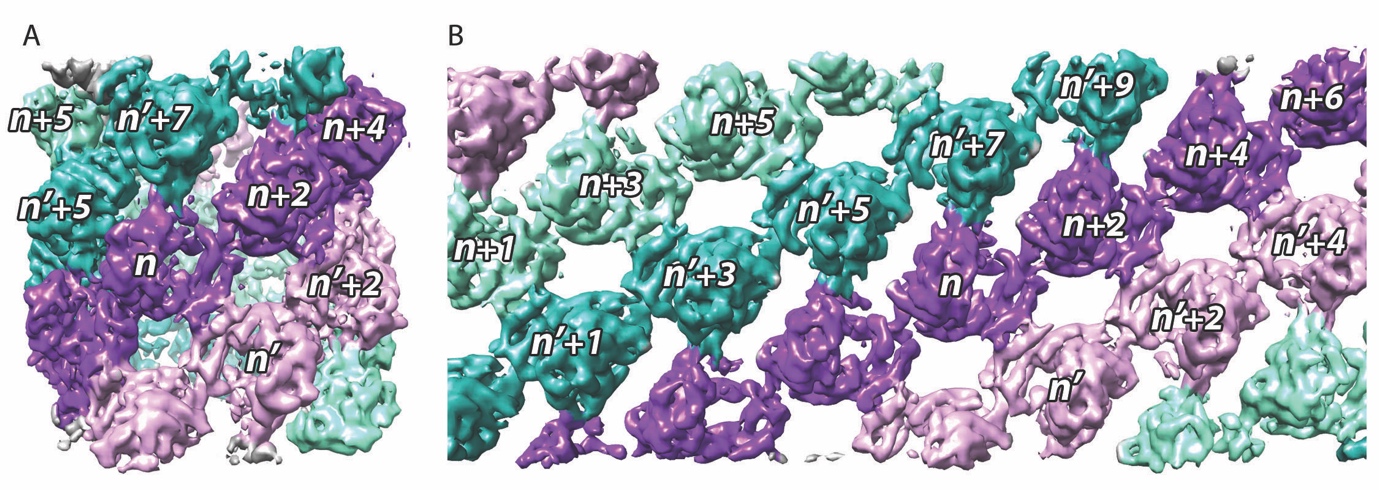

**Figure S4**

The cryo-EM reconstruction for the tight/narrow fibril conformation N3 is shown coloured to highlight the two protofilaments present. Protofilament 1 is shown as purple/plum, while protofilament 2 is shown as aquamarine/light sea green. The notation used to describe interactions between 3D^pol^ is illustrated both on the reconstructed density map (A), and on a representation of the same map that has been unrolled to produce a planar array (B).

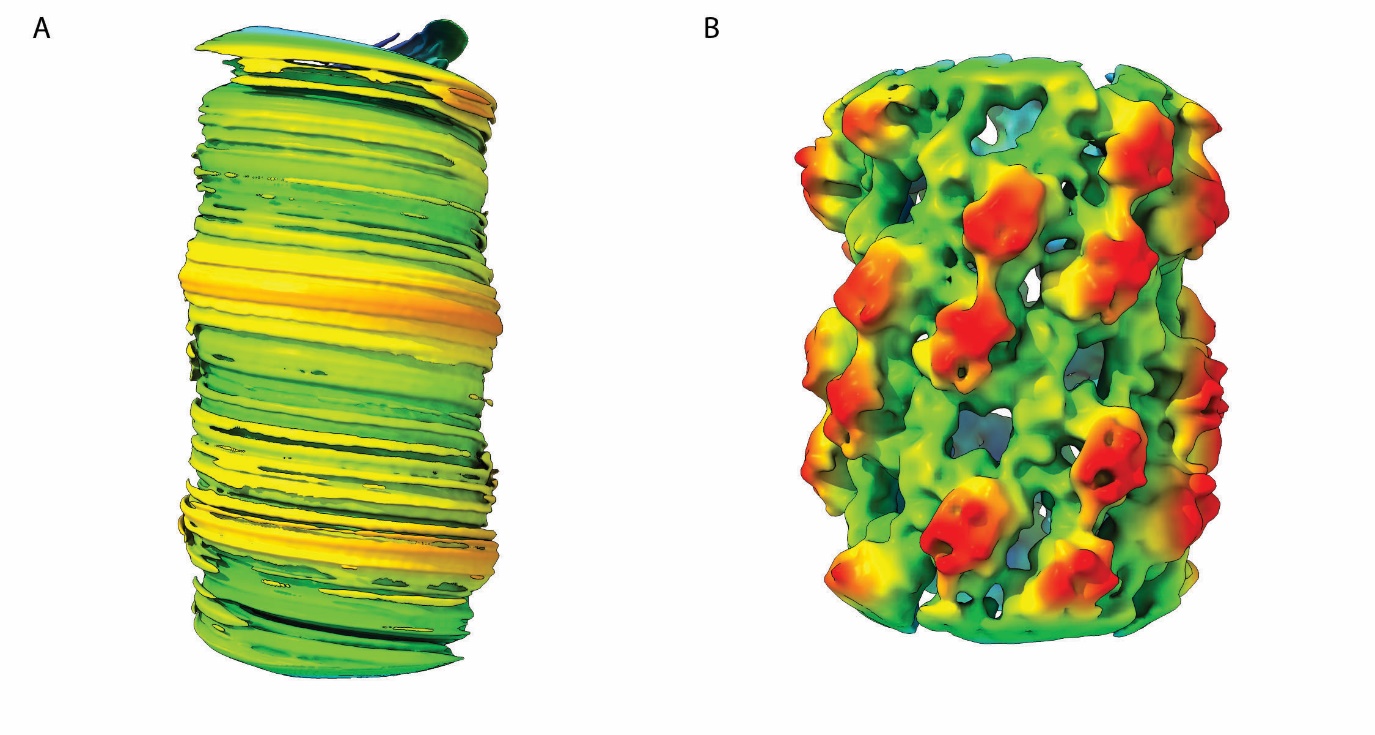

**Figure S5**

Preliminary 3D reconstruction analysis of 3D^pol^ fibrils. A small dataset was collected on a JEOL 2200 cryomicroscope equipped with a Direct Electron DE20. An initial model was generated (A) and used as a starting point for helical reconstruction, which led to an interpretable density map (B).

**Supplemental Table 1**

Table to show imaging and reconstruction statistics for each of the fibril reconstructions.

**Supplemental Table 2**

Tables to show amino-acid residues identified in contacts/clash analyses of interfaces (A) between monomers in a single dimeric subunit (dimer interface), (B) between dimers leading to the formation of ribbons of dimers (protofilaments) and (C) between protofilaments leading to the formation of fibrils. Contact residues common to all dimer and dimer-dimer interfaces are coloured red. Inter-protofilament contact residues common to tight forms are coloured orange, those common to open forms are coloured blue.

**Supplemental Movie 1**

Movie to show nine fibril conformations determined by cryo-EM and helical reconstruction. Each conformation is shown as a radially coloured isosurface. Conformation N3 is shown as a transparent isosurface with a monomer of 3D^pol^ as a rainbow coloured ribbon diagram (timepoint 2.02). A second 3D^pol^ molecule is shown to illustrate the formation of dimers (timepoint 2.17). Protofilament one is shown as a ribbon of dimers coloured violet and thistle (timepoint 2.34). Protofilament two is shown coloured teal and aquamarine (timepoint 2.42).

Morphs are presented between conformations using solvent excluded surface representations. Conformations N3 to N8 show similar packing of protofilaments. Variation in fibril diameter is a result of protofilaments flexing/sliding. Narrow conformations have 8.4-8.57 dimers in each protofilament per helix turn. Morph of N3-N8 highlights the closing together of protofilaments resolved for the tight/narrow forms (timepoint 2.48 to 2.54). Conformation B1 also shows protofilaments packing tightly together, however in this conformation the two protofilaments are symmetric across the helix axis (C2 symmetry). Broad conformations have 9-9.2 dimers per turn in each protofilament. Morph of B1-N3 illustrate the difference between tight/narrow and tight/broad conformations (timepoint 2.58-3.02). For narrow and broad fibrils, reconstructions were calculated that showed a more open conformation - B2 and N9. Similar to the densely packed fibrils, these structures were formed of protofilaments of dimers. The contact interface between protofilaments was rather different however. Morph animations are intended to highlight how these open conformations differ from densely packed fibrils, showing how protofilaments slide against each other leading to opening of a cavity between the two ribbons and a further narrowing of the fibril (timepoints 1.45 to 1.50). Morphs between tight and open conformations (B1-B2) timepoint 3.06-3.11 (N3-N9) timepoint 3.16-3.23.

**Supplemental Movie 2**

Movie to show protein-protein interactions that lead to the formation of FMDV 3D^pol^ fibrils. N3 is shown as a transparent isosurface with a pseudoatomic model of the fibril shown coloured to highlight the domain structure of the RNA dependent RNA polymerase; palm domain (plum), fingers domain (powder blue) and thumb domain (gold). Contacts/clashes at the dimer interface (timepoint 0.37), inter-dimer interface (timepoint 1.00) and inter-protofilament interface (timepoint 1.24) are shown as white dashed lines. Fibril N9 is shown as a transparent isosurface with fitted atomic models (timepoint 1.37), the inter-protofilament contact interface is highlighted (timepoint 2.08). The interface between protofilaments may involve the formation of an extended β-sheet, putative hydrogen bonds are indicated by blue dashed lines.

**Supplemental Movie 3**

Movie to show cryo-EM density that is not assigned to protein by rigid-body fitting of PDB 2EC0, we hypothesise that density seen in the active site of 3D^pol^ indicates the presence of RNA within fibrils that may be at low-occupancy or poorly ordered; N3 is shown as a radial-coloured isosurface cryo-EM map. Putative RNA density in the active site is indicated by a white arrow (timepoint 0.28). A transparent map is shown with fitted atomic coordinates for both protein and nucleic acid, as determined by (38 – timepoint 0.34). The open fibril form N9 has a large finger of density that extrudes from the active site and extends between protofilaments. We interpret this as being RNA (timepoint 0.57).
